# Navigating Chemical Space By Interfacing Generative Artificial Intelligence and Molecular Docking

**DOI:** 10.1101/2020.06.09.143289

**Authors:** Ziqiao Xu, Orrette Wauchope, Aaron T. Frank

**Affiliations:** the University of Michigan, Department of Chemistry, Ann Arbor, 48109, United States of America; City University of New York, Baruch College, Department of Natural Sciences, New York, 10010, United States of America; the University of Michigan, Biophysics, Ann Arbor, 48109, United States of America

**Keywords:** Virtual Screening Libraries, Biophysical Modeling, Computational Drug Discovery

## Abstract

Here we report the testing and application of a simple, structure-aware framework to design target-specific screening libraries for drug development. Our approach combines advances in generative artificial intelligence (AI) with conventional molecular docking to rapidly explore chemical space *conditioned* on the unique physiochemical properties of the active site of a biomolecular target. As a proof-of-concept, we used our framework to construct a focused library for cyclin-dependent kinase type-2 (CDK2). We then used it to rapidly generate a library specific to the active site of the main protease (M^pro^) of the SARS-CoV-2 virus, which causes COVID-19. By comparing approved and experimental drugs to compounds in our library, we also identified six drugs, namely, Naratriptan, Etryptamine, Panobinostat, Procainamide, Sertraline, and Lidamidine, as possible SARS-CoV-2 M^pro^ targeting compounds and, as such, potential drug repurposing candidates. To complement the open-science COVID-19 drug discovery initiatives, we make our SARS-CoV-2 M^pro^ library fully accessible to the research community (https://github.com/atfrank/SARS-CoV-2).

---

Developing new drugs is costly and time-consuming; by some estimates, bringing a new drug to market requires ~13 years at the cost of ~US$1 billion.(1) As such, there is a keen interest in accelerating the drug development pipeline and reduce its cost. In principle, computer-aided drug development can both accelerate and lower the cost of identifying viable drug candidates.(2) For instance, during hit identification, where the goal is to identify small molecules that may bind to the target, one could restrict testing to only the compounds that are predicted to bind to the target with high affinity. However, the computational cost of screening large chemical libraries can still be burdensome. The computation cost of virtual screening could be reduced by working with smaller and more focused, target-specific virtual libraries, enriched in compounds that are likely to bind to a specific site on the target of interest.(3) Such libraries can be constructed by carrying out *de novo* design, during which compounds are designed on-the-fly, guided by the unique physiochemical properties of the active site of the target.(4–17) However, because of the difficulty in formalizing diverse reaction rules and implementing robust algorithms to intelligently apply them, up to this point, such *de novo* design frameworks often produce synthetically inaccessible compounds.

Fortunately, advances in generative artificial intelligence now make it possible to design both chemically novel and synthetically feasible compounds in silico.(18–23) Noteworthy among these frameworks are those based on autoencoders. Autoencoders are unsupervised machine learning models that, by virtue of their architecture, are able to learn a compact, latent space representation of chemical space when they are trained with molecular data. The task of designing target-specific libraries can, therefore, be cast as one of exploring the latent space of such models, conditioned on the unique properties of the active site on the target of interest. Currently lacking, however, are efficient, structure-aware strategies for sampling the latent space of generative molecular models. If developed, such techniques could find utility in constructing active-site and target-specific virtual libraries that are likely to contain hit compounds that, if synthesized, might bind to the targeted site. More immediately, they could be used to construct a ligand-based pharmacophore model that can be used to screen millions of compounds in a matter of seconds.(24)

Here, we tested and implemented a novel, structure-based framework for constructing target-specific libraries by sampling the latent space of generative machine learning models *conditioned* on the unique physiochemical properties of the “active site” of a target. We reasoned that we could implement such a pipeline by combining generative molecular machine learning models with molecular computer docking algorithms. For instance, one could start by embedding a reference compound, e.g., benzene, into the latent space of a generative autoencoder model and sample latent space points in its “neighborhood.” The sampled points can then be decoded into their corresponding 3-dimensional (3D) representations and then docked onto the target (Figure 1a). Of these sampled compounds, the one that is predicted to interact most favorably with the active site can be identified and then used as the new reference. The process can be repeated (Figure 1b), and in so doing, the latent-space of a generative molecular model (and, so chemical space) explored in a semi-directed manner, *conditioned* on the properties of the active site of the target. The compounds sampled during this iterative (“sample-and-dock”) process could then be used to construct a target-specific virtual library.

**Fig. 1.**
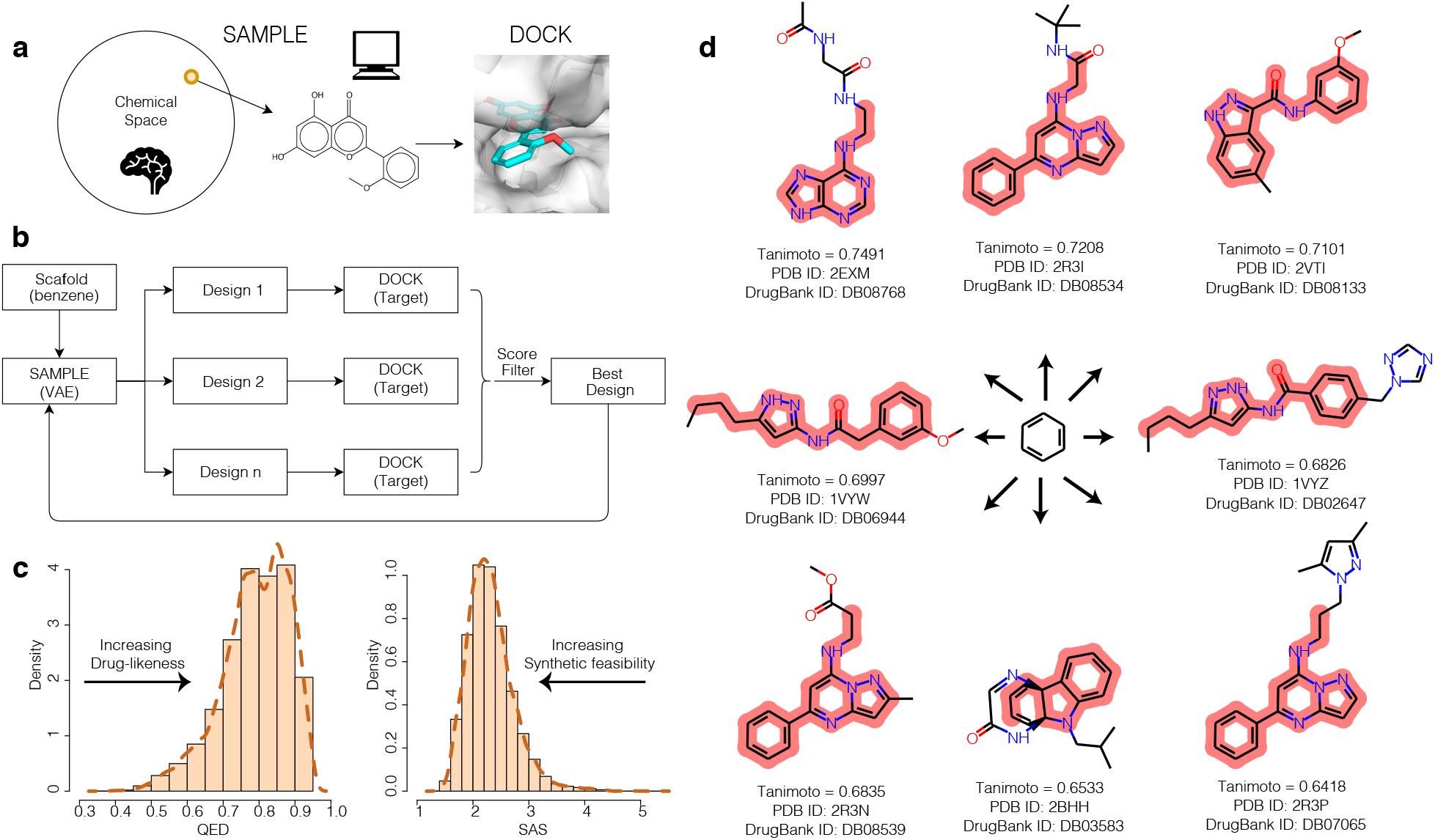
Illustration of (a) the “sample-and-dock” concept that can be used to implement (b) a structure-aware design pipeline. (c) Computed properties of compounds design to target the active site of CDK2. (d) Representative compounds in our sample-and-dock library that closely resemble known inhibitors of CDK2. Highlighted are sub-substructures that are identical to those found in the most similar compound in our sample-and-dock virtual library.

First, we implemented a sample-and-dock pipeline and then used it to construct a target-specific library for the cyclin-dependent kinase 2 (CDK2) protein. We chose CDK2 because it is an important therapeutic target that has been extensively studied biochemically and structurally, and for which a large library of known inhibitors is readily accessible.(25) To achieve this, we first interfaced the junction-tree variational autoencoder (JTVAE)(26) with the docking program, rDock.(27) JTVAE was chosen because it has demonstrated a superior ability to generate synthetically feasible molecules, and rDock was chosen because it is fast, and the software is open-sourced. We then applied our JTVAE-rDock sample-and-dock pipeline (Figure 1a and b) to CDK2. For this CDK2 test case, we initiated sample-the-dock pipeline by first mapping the CDK2 active site using the crystal structure of CDK2 complexed with Roniciclib (PDB: 5IEV).(28) The JTVAE-rDock sample-and-dock pipeline was then initialized using benzene as a starting scaffold. We chose to use benzene as the starting scaffold because it is small and a chemical motif that is ubiquitous in known drugs. Each sample-and-dock cycle consisted of sampling and docking 20, unique compounds. For each compound, 100 poses were generated during docking. The compound with the highest predicted binding affinity was selected and used as the input scaffold for the next cycle of sample-and-dock.

To explore the quality of the compounds sampled during sample-and-dock, we computed the distribution of their drug-likeness (estimated using the quantitative estimate of drug-likeness, QED,(29) score) and synthetic feasibility (estimated using their synthetic accessibility score, SAS). Though there are exceptions, compounds suitable for initializing drug-development typically exhibit QED and SAS →1.0, respectively. For CDK2, the sample-and-dock molecules exhibited mean QED and SAS values of ~0.80 and 2.5, respectively (Figure 1c). To examine whether the compounds explored during sample- and-dock resemble known inhibitors of CDK2, we computed their chemical similarity to known CDK2 inhibitors. Briefly, we identified known CDK2 inhibitors within the DrugBank,(25) and calculated the Tanimoto coefficient between the chemical fingerprints of the known inhibitors and the sample-and-dock compounds. Remarkably, despite initializing sample-and-dock with just benzene, we found that we were indeed able to sample designs that closely resemble known CDK2 inhibitors (Figure 1d). Collectively, therefore, our results for CDK2 suggest that starting from available structure and knowledge of the binding site, generative machine learning models and molecular docking could be seamlessly interfaced to rapidly construct a target-specific screening library comprised of compounds that are predicted to be both drug-like and synthetically feasible and are likely to resemble known small molecule modulators of the target.

Next, we applied our sample-and-dock framework to the main protease (M^pro^) of the virus, SARS-CoV-2. SARS-CoV-2 M^pro^ processes polypeptides that are important for viral assembly and, as such, is an attractive target for developing drugs to treat COVID-19.(30, 31) To generate a SARS-CoV-2 M^pro^-specific library, we initiated sample-and-dock calculations starting with benzene as the reference scaffold and targeting the same site on SARS-CoV-2 M^pro^ that is occupied by the inhibitor, N3 (PDB: 6Y2F).(32) The active site of SARS-CoV-2 M^pro^ is comprised of five sub-pockets, which we denote as P-I, P-II, P-III, P-IV, and P-V (Figure 2a). Over 48 hours, we generated a virtual library containing 1448 compounds. Shown in Figure 2 are three representative compounds within our SARS-CoV-2 M^pro^ library. Inspection of the predicted poses indicate that the three representative compounds occupied pockets P-I, P-II, and P-IV (Figure 2a), and could form stabilizing contacts with the catalytic residues, His^41^ and Cys^145^ (Figure 2b) Interestingly, both SD-2020-1 (QED=0.51) and SD-2020-2 (QED=0.59) contain the peptidyl moiety found in known M^pro^ inhibitors(30, 32). In fact, the presence of the peptidyl moiety was a common feature of many of the compounds that were sampled during sample-and-dock. On the other hand, SD-2020-3 (QED=0.75), a nucleotide-analog, is an example of one of the compounds in our library that lacks the peptidyl moiety. The overall drug-likeness, and lack of the reactive peptidyl moiety, make SD-2020-3 an interesting lead candidate. On the basis of the results for CDK2, we speculate that the SARS-CoV-2 M^pro^ specific library we generated using our sample-and-dock approach may contain yet to be explored and promising lead candidates. Accordingly, we make the library of M^pro^ designs fully open to the scientific community with the hope that it will be explored by medicinal chemists to seed the design of novel drugs targeting M^pro^ of SAR-CoV-2 (https://github.com/atfrank/SARS-CoV-2).

**Fig. 2.**
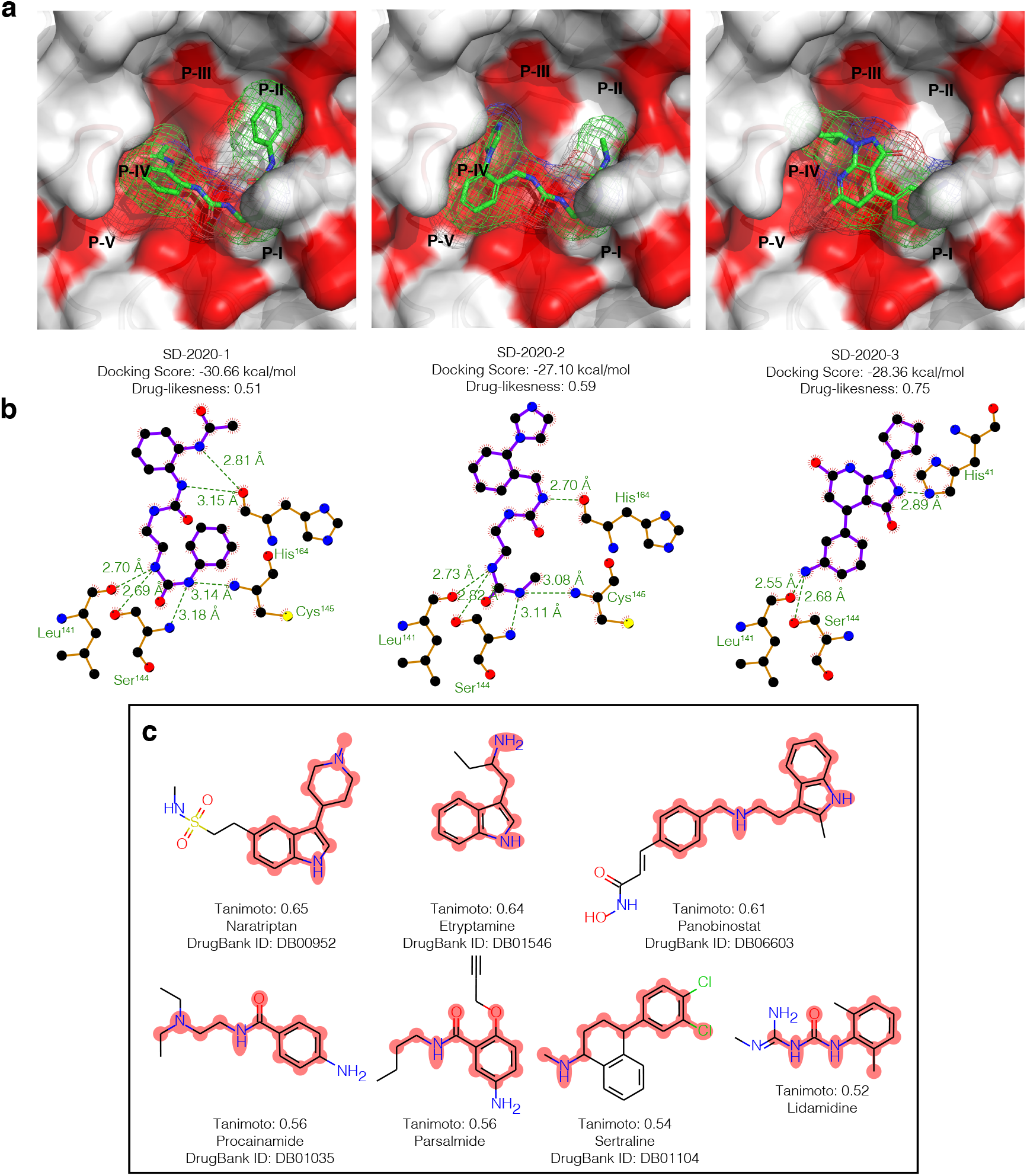
(a) Examples of three potential inhibitors (SD-2020-1, −2, and −3) designed using the sample-and-dock approach. Shown are the predicted poses of these compounds, docked into the active site of SARS-CoV-2 M^pro^. (b) Ligand interaction maps of the docked poses. Docking predicts that the SD-2020-1 and SD-2020-3 interact with the catalytic residue Cys^145^, and SD-2020-2 interacts with the catalytic residue His^41^. (c) Known drugs that were most similar to compounds in our sample-and-docking M^pro^ library. Highlighted are sub-substructures that are identical to those found in the most similar compound in the virtual library.

Finally, we asked whether the compounds in our virtual SAR-CoV-2 M^pro^ library were similar to any known or experimental drugs; identifying such drugs would be one way we could immediately leverage our virtual library to assist ongoing COVID-19 drug repurposing efforts. To answer this question, we compared the compounds in our library with compounds in SuperDRUG2,(33) a library comprised of FDA approved and experimental drugs. Shown in Figure 2c are the six drugs that exhibited the highest chemical similarity to at least one compound in our SAR-CoV-2 M^pro^ library. Of these six, three are central nervous system drugs, which include Sertraline, a selective serotonin reuptake inhibitor (SSRI). Intriguingly, another SSRI, fluvoxamine, has recently been identified as a potential COVID-19 drug and is now in clinical trials.

To summarize, we combined emerging generative modeling with well-established biophysical modeling to yield a hybrid approach for the structure-guided exploration of chemical space. Instead of exploring chemical space in a large and fixed library, our approach facilitates a directed exploration of the chemical space that is relevant to a specific active site on a given target. The sample-and-dock pipeline we implemented is also modular and flexible. For instance, as more robust generative models are developed, they can be trivially incorporated into our pipeline, as can more accurate and robust scoring functions and methods. In its current form, however, our sample-and-dock pipeline can be deployed to construct structure- and target-specific maps of the “privileged” chemical space for therapeutically relevant protein and nucleic acid targets. Applying unsupervised learning to sets of structure- and target-specific chemical-space maps could facilitate the emergence of general design principles that can guide the rational structure-guided design of biomolecular probe compounds.(34) Accordingly, we have implemented the sample-and-dock pipeline into an easy-to-use software we refer to as SampleDock and make it accessible to the research community at (https://smaltr.org/).

## Supporting Information (SI)

### Software

The SampleDock software can be access at https://smaltr.org/.

### Data sets

The sample-and-dock library for SARS-CoV-2 M^pro^ (as well as other COVID-19 target) can be access at https://github.com/atfrank/SARS-CoV-2.

## Materials and Methods

### Sample-and-Dock

To construct structure-aware and target-specific virtual libraries, we implemented a *conditional* generative sampling scheme. Our *conditional* generative sampling scheme, which we refer to as sample-and-dock, combines a pre-trained generative variational autoencoder (VAE) model with molecular docking (see below). In sample-and-dock, the exploration of the latent space of a pre-trained generative model is made *conditional* on the unique physiochemical and biophysical properties of the binding site on a target, 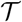, by using a greedy algorithm that optimizes a cost function that dependent of the properties of the target. Here we use molecular docking scoring function as the cost function in our greedy algorithm. Accordingly, given a set of latent-space samples, the “best” is selected as the one that, when decoded into its molecular representation, yields the highest binding affinities 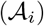, as estimated using a molecular docking scoring function.

Briefly, to construct a customized target-specific library using this conditional sampling scheme, we first embed a reference compound in the latent space of a generative model (*m_i_ → z_i_*) and then **sample** a set of {*z*_*i*+1_} in the “neighborhood” of the reference *z_i_*. The {*z*_*i*+1_} can then be decoded (*z*_*i*+1_ → *m*_*i*+1,*D*_) and the resulting molecules **docked** into the active site of 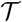. Using a greedy optimization strategy, the “best” of these designed molecules can be identified using by the docking-derived 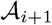 and then it is used as the new reference for the next round of **sample** and **dock**. The process is then repeated over many cycles to explore the chemical space in a semi-directed manner and *conditioned* on the properties of 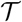.

### Implementation

Using a set of in-house python scripts, we implemented a sample-and-dock pipeline that interfaced with the junction-tree-variation autoencoder (JT-VAE) model of Jin and coworkers,(26) with the molecular docking program rDock.(27). All instances of sample-and-dock were initialized using benzene. At the start of each cycle, the seeding molecule was taken in the form of SMILES strings and converted into one-hot encoding according to a set of predefined vocabulary. The one-hot encoding was first transformed into a set of two 28-dimensional vectors by the encoders of JTVAE as locations in the two latent spaces. The latent vectors were then re-sampled and reconstructed (decoded to SMILES strings) 20 times by the decoders to generate 20 unique molecules that resemble the seed. For each generated molecule, the SMILES string was converted to a 3D structure using RDKit and then docked into the active site on the target using rDock. Across all the generated molecules within the cycle, the designed molecule with the highest 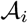, as estimated using the rDock scoring function, was used to as the seed molecule for the next cycle. The process is repeated over many cycles, and the best compound from each cycle used to construct the final library.

### Target-Specific Libraries

We used our sample-and-dock pipeline to construct target-specific libraries for CDK2 and SAR-CoV-2 M^pro^. For CDK2, sample-and-dock was carried out using the crystal structure of CDK2 bound with Roniciclib (PDB ID: 5IEV).(28) For SAR-CoV-2 M^pro^, we used the crystal structure M^pro^ bound to N3 (PDB ID: 6Y2F).(32) From the crystal structures, the ligands and receptors separated and saved as separate coordinate files using PyMOL. To map the active sites of each target, cavity mapping was carried out using the reference-ligand method that is implemented in the rbcavity tool from rDock. For the reference-ligand method, the radius of the overlapping sphere was set to 7.0 Å, and the radius of the small probe radii was set to 1.5 Å. For both CDK2 and M^pro^, sample-and-dock was initialized using benzene and ran for 48 hours.

## Supporting Information

**Fig. S1.**
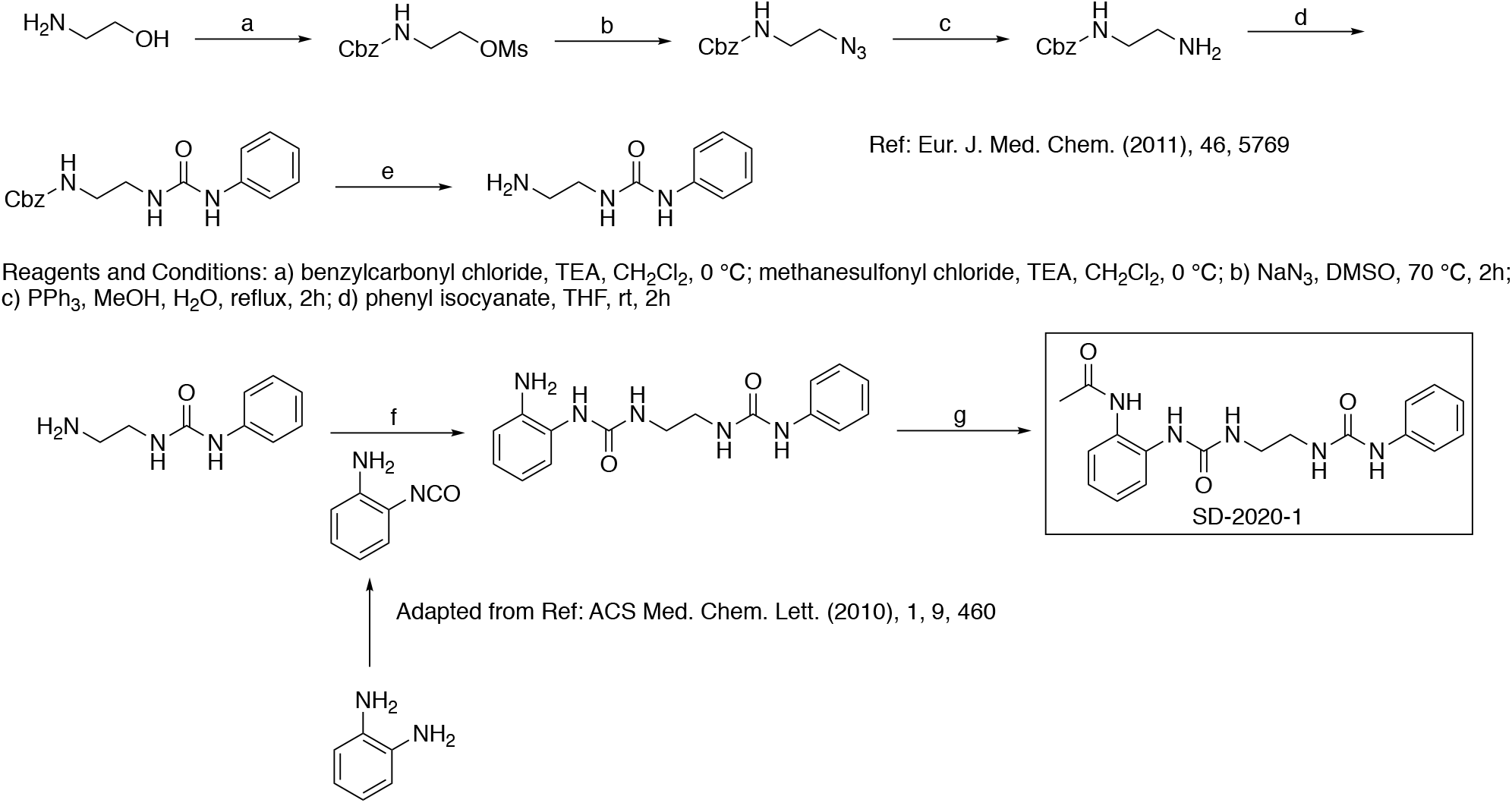
Proposed synthetic scheme for SD-2020-1.

**Fig. S2.**
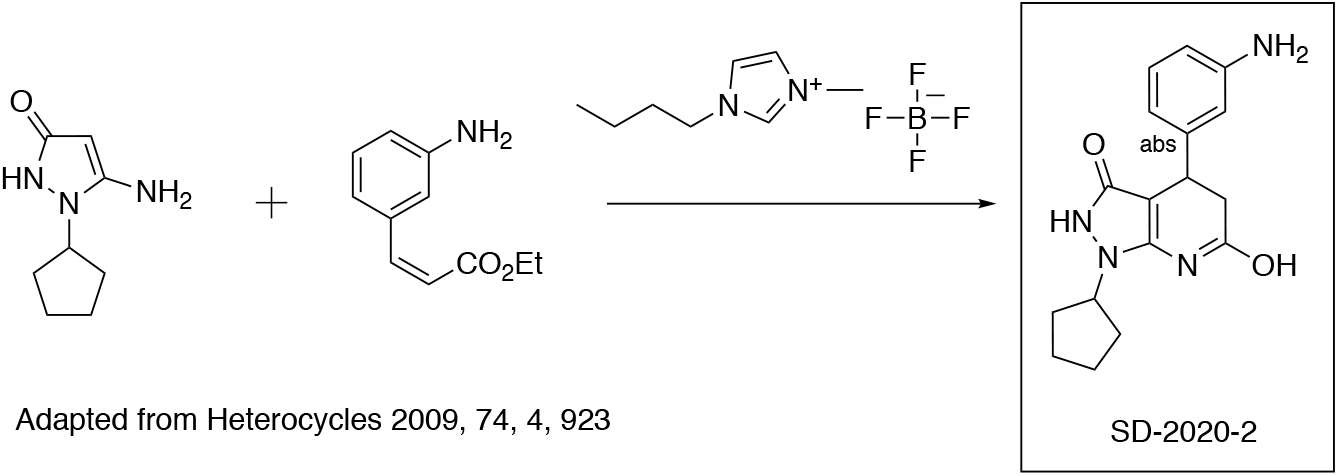
Proposed synthetic scheme for SD-2020-2.

**Fig. S3.**
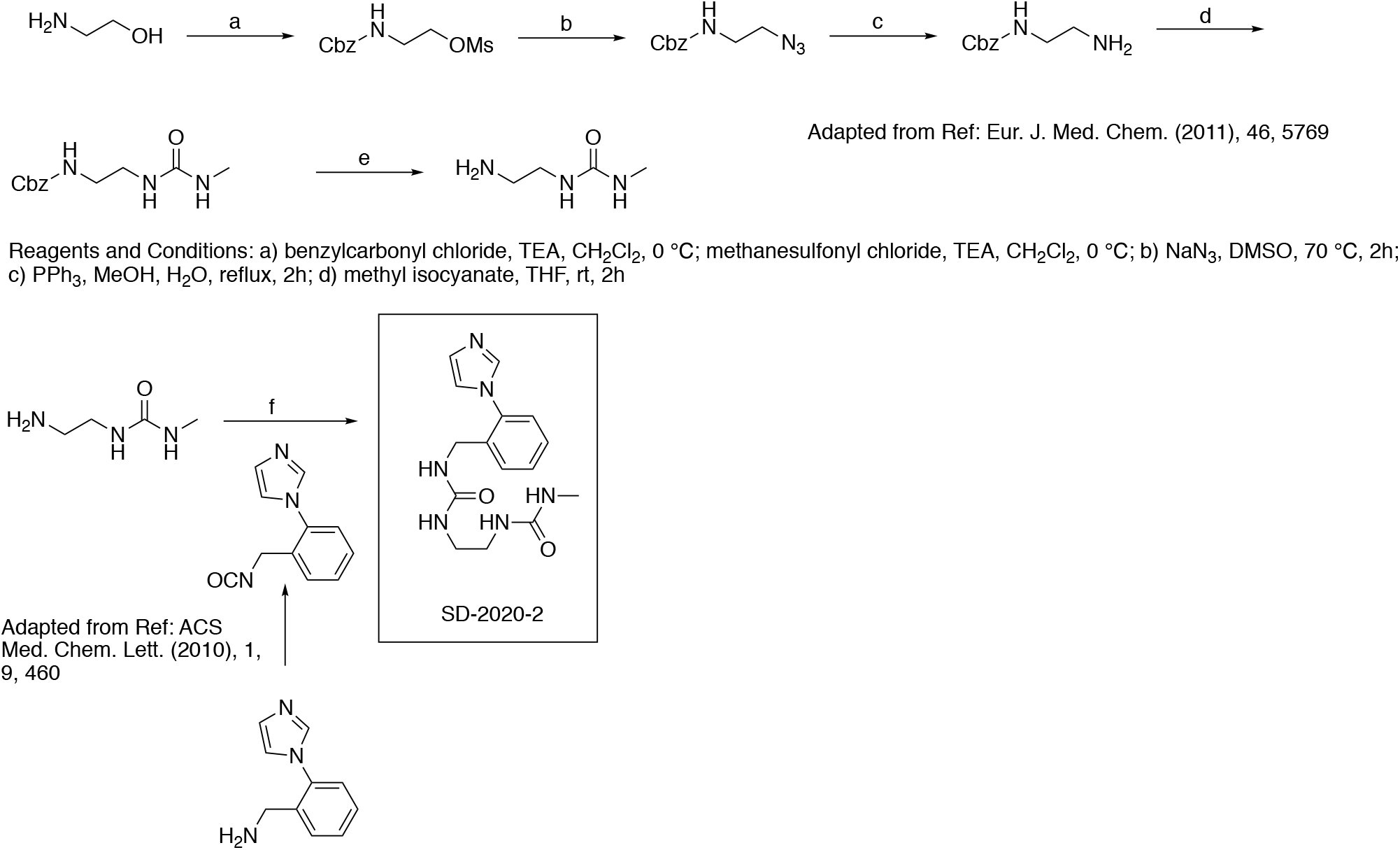
Proposed synthetic scheme for SD-2020-3.

## Notes

https://github.com/atfrank/SARS-CoV-2

### Competing Interest Statement

The authors have declared no competing interest.

